# Biotic and abiotic drivers of soil microbial functions across tree diversity experiments

**DOI:** 10.1101/2020.01.30.927277

**Authors:** Simone Cesarz, Dylan Craven, Harald Auge, Helge Bruelheide, Bastien Castagneyrol, Andy Hector, Hervé Jactel, Julia Koricheva, Christian Messier, Bart Muys, Michael J. O’Brien, Alain Paquette, Quentin Ponette, Catherine Potvin, Peter B. Reich, Michael Scherer-Lorenzen, Andrew R Smith, Kris Verheyen, Nico Eisenhauer

**Affiliations:** German Centre for Integrative Biodiversity Research (iDiv) Halle-Jena-Leipzig, Deutscher Platz 5e, 04103 Leipzig, Germany; Institute of Biology, Leipzig University, Johannisallee 21, 04103 Leipzig, Germany; Institute of Ecology, Friedrich-Schiller-University Jena, Dornburger Str. 159, 07749 Jena, Germany; Centro de Modelación y Monitoreo de Ecosistemas, Facultad de Ciencias, Universidad Mayor, Santiago, Chile; Department of Community Ecology, Helmholtz‐Centre for Environmental Research – UFZ, Theodor-Lieser-Str. 4, 06120 Halle, Germany; Martin Luther University Halle-Wittenberg, Institute of Biology/Geobotany and Botanical Garden, Am Kirchtor 1, 06108 Halle, Germany; INRAE (The French National Institute for Agriculture, Food and Environment), Biogeco, 69 Route d’Arcachon, 33612 Cestas, France; Department of Plant Sciences, University of Oxford, OX1 3RB, UK; Department of Biological Sciences, Royal Holloway University of London, Egham, Surrey, TW20 0EX, UK; Département des Sciences Naturelles, Université du Québec en Outaouais (UQO), ISFORT, Ripon, Canada; Centre for forest research, Université du Québec à Montréal, PO Box 8888, Centre-ville station, Montréal, Qc, Canada H3C 3P8; Division Forest, Nature & Landscape KU Leuven, Celestijnenlaan 200E box 2411, 3001 Leuven, Belgium; Área de Biodiversidad y Conservación, Universidad Rey Juan Carlos, c/Tulipán s/n., E-28933 Móstoles, Spain; Earth & Life Institute, Université catholique de Louvain (UCLouvain), Croix du Sud 2 - box L7.05.09, 1348 Louvain-la-Neuve, Belgium; Department of Biology, McGill university, Montreal, Canada; Department of Forest Resources, University of Minnesota, St. Paul, MN 55108, USA; Hawkesbury Institute for the Environment, Western Sydney University,Penrith, NSW 2751, Australia; Geobotany, Faculty of Biology, University of Freiburg, Schänzlestr. 1, 79104 Freiburg, Germany; School of Natural Sciences, Bangor University, Bangor, Gwynedd, LL57 2UW, UK; Forest & Nature Lab, Department of Environment, Ghent University, Geraardsbergsesteenweg 267, B-9090 Melle-Gontrode, Belgium

**Keywords:** Aboveground-belowground interactions, Biodiversity-ecosystem functioning, Soil biota, Soil microorganisms, Biodiversity loss, TreeDivNet

## Abstract

**Aim:** Soil microorganisms are essential for the functioning of terrestrial ecosystems. Although soil microbial communities and functions may be linked to the tree species composition and diversity of forests, there has been no comprehensive study of how general potential relationships are and if these are context-dependent. A global network of tree diversity experiments (TreeDivNet) allows for a first examination of tree diversity-soil microbial function relationships across environmental gradients.

**Location:** Global

**Major Taxa Studied:** Soil microorganisms

**Methods:** Soil samples collected from eleven tree diversity experiments in four biomes across four continents were used to measure soil basal respiration, microbial biomass, and carbon use efficiency using the substrate-induced respiration method. All samples were measured using the same analytical device in the same laboratory to prevent measurement bias. We used linear mixed-effects models to examine the effects of tree species diversity, environmental conditions, and their interactions on soil microbial functions.

**Results:** Across biodiversity experiments, abiotic drivers, mainly soil water content, significantly increased soil microbial functions. Potential evapotranspiration (PET) increased, whereas soil C-to-N ratio (CN) decreased soil microbial functions under dry soil conditions, but high soil water content reduced the importance of other abiotic drivers. Tree species richness and phylogenetic diversity had overall similar, but weak and context-dependent (climate, soil abiotic variables) effects on soil microbial respiration. Positive tree diversity effects on soil microbial respiration were most pronounced at low PET, low soil CN, and high tree density. Soil microbial functions increased with the age of the experiment.

**Main conclusions:** Our results point at the importance of soil water content for maintaining high levels of soil microbial functions and modulating effects of other environmental drivers. Moreover, overall tree diversity effects on soil microbial functions seem to be negligible in the short term (experiments were 1-18 years old). However, context-dependent tree diversity effects (climate, soil abiotic variables) have greater importance at high tree density, and significant effects of experimental age call for longer-term studies. Such systematic insights are key to better integrate soil carbon dynamics into the management of afforestation projects across environmental contexts, as today’s reforestation efforts remain focused largely on aboveground carbon storage and are still dominated by less diverse forests stands of commercial species.

## Introduction

Soil microorganisms are the functional backbones of terrestrial ecosystems (van der Heijden et al. 2008) as they underpin crucial ecosystem functions and services that humankind relies on (Wall et al., 2015). Given the critical role of soil microorganisms in carbon dynamics (Gougoulias et al., 2014) and soil feedback effects on climate (Classen et al., 2015), improving current understanding of the drivers of microbial biomass and activity is an essential step towards predicting global change impacts (Serna-Chavez et al., 2013; Xu et al., 2013; Thakur et al., 2015). Soil microbial biomass can serve as a proxy for nutrient cycling and soil enzyme dynamics, such as soil organic matter (SOM) turnover as well as for secondary productivity (Crowther et al., 2019). In addition, in-situ measurements of microbial activity have been correlated to rates of soil C sequestration (Lange et al., 2015). Together, microbial biomass and microbial activity provide critical information on a range of important soil ecosystem functions (Eisenhauer et al., 2018).

Globally, abiotic factors are thought to drive variation in soil microbial biomass and microbial activity (Fierer et al., 2009; Serna-Chavez et al., 2013). High soil moisture, neutral soil pH, low soil C/N ratio, and high soil organic carbon content (here summarized as high soil quality) are among the most important factors directly increasing soil microbial biomass and activity. In contrast, climatic conditions like temperature may influence soil microbial biomass indirectly by soil water content *via* evapotranspiration and changes in soil organic matter content (Fierer et al., 2009; Serna-Chavez et al., 2013). These patterns become less clear when taking into account interactions among different drivers. For instance, positive effects of high soil nutrient content can be constrained by stressful environments, e.g., in dry and nutrient limited systems (Strickland & Rousk, 2010; Serna-Chavez et al., 2013), highlighting the importance of context-dependent effects. Moreover, effects of abiotic drivers may further be modulated by local biotic conditions. For example, recent studies in grasslands have demonstrated that plant diversity affects soil microbial community composition, activity, and biomass (Lange et al., 2015; Thakur et al., 2015) with significant effects on ecosystem functions, such as soil carbon storage (Lange et al., 2015), litter mineralization (Pei et al., 2017), and soil N retention (Leimer et al., 2016). However, existing global analyses of plant diversity effects on soil microbial communities have revealed inconsistent results and focused either on soil communities, but not on soil functions, or grasslands only (Tedersoo et al., 2014, 2016; Prober et al., 2015; Thakur et al., 2015). Inconsistencies in the magnitude and direction of biodiversity effects may be due to strengthening biodiversity effects with time and different environmental contexts, such as different soil conditions (Guerrero-Ramírez et al., 2017). So far, plant diversity effects on soil microbial functions have been studied mostly in grasslands, and little is known for forests (Chen et al., 2019). This is a major knowledge gap, because there might be substantial differences between ecosystems in terms of soil microbial function and potential climate feedback effects on soil communities (Chen et al., 2018).

Previous studies on tree diversity effects on soil microorganisms mainly compared monoculture stands with mixtures of two tree species in different environments, making it difficult to disentangle site conditions from tree diversity and tree identity effects (Scheibe et al., 2015; Chodak et al., 2016; Crowther et al., 2016). One of the first studies using data from a tree diversity experiment with homogeneous abiotic conditions found soil microbial activity and biomass to increase with tree species richness in a saturating relationship, while soil microbial community composition did not vary significantly (Khlifa et al., 2017). One of the potential mechanisms underlying a positive plant diversity effect on microorganisms is the increased input of diverse resources (Steinauer et al., 2016). In line with the view that the quality of plant inputs are essential for soil microbial processes, the chemical composition of leaf litter determines nutrient mineralization, microbial respiration, and microbial biomass (Pei et al., 2016), whereas species diversity *per se* was shown to have little effect (Meier & Bowman, 2008; Steinauer et al., 2016). This finding suggests that an increase in species richness may not increase soil microbial biomass and activity if not accompanied with a simultaneous increase in functional dissimilarity of co-occurring species (Heemsbergen et al., 2004). Although debated, research in grasslands suggests that functional diversity is of higher importance than plant diversity (e.g., Flynn et al., 2011), while there is even less conclusive information from forest ecosystems (Scherer-Lorenzen et al., 2007). Unfortunately, access to and measuring the same belowground traits (e.g., leaf N concentration, leaf lignin concentration, mycorrhizal association) is often not possible for logistical reasons. To overcome this lack of data, phylogenetic diversity can be used as a proxy for functional diversity (Tucker et al., 2018). In addition, long-term diversity experiments show that biodiversity effects strengthen with time. Suggested mechanisms are an increase in the complementarity due to increased functional diversity and reduced redundancy of species in a community (Reich et al., 2012; Strecker et al., 2016; Guerrero-Ramírez et al., 2017). Such context-dependent plant diversity effects call for systematic studies across diversity gradients and environmental contexts.

Here, we present the first coordinated analysis of data on soil microbial functions related to soil carbon dynamics across eleven tree diversity experiments distributed across four biomes (Paquette et al., 2018). To explore potential tree diversity effects on soil microbial functions, we tested tree species richness (the biodiversity measure most frequently manipulated in tree diversity experiments; Verheyen et al., 2016) and tree phylogenetic diversity effects. We expect that phylogenetically diverse tree stands will provide more dissimilar resources and niches to soil microorganisms, thereby increasing ecosystem functioning. Specifically, we tested the responses of three soil microbial functions. First, soil microbial basal respiration reflects soil microbial activity, which provides an activity measure at the moment of sampling as only a fraction of the community is active due to environmental constraints (e.g. soil moisture, nutrient availability). Second, soil microbial biomass was measured as a proxy for secondary productivity (i.e., production of biomass of heterotrophic organisms). Third, soil microbial specific respiratory quotient was used as an indicator of microbial carbon use efficiency and is the ratio of basal respiration and microbial biomass. Our main hypotheses were (H1) that tree diversity will increase soil microbial biomass, activity, and carbon use efficiency (Chen et al., 2019), and (H2) that these effects will be stronger in tree mixtures with a higher phylogenetic diversity, reflecting greater differences in a range of tree traits (Cadotte et al., 2009). In addition, we expected abiotic drivers to strongly influence soil microbial functions. High soil carbon (soil C) concentration, high soil moisture, and neutral soil pH were hypothesized to increase the biomass, activity, and carbon use efficiency of soil microorganisms (H3). However, abiotic and biotic drivers may interact in influencing soil microbial functions given the context dependency of biodiversity-ecosystem function relationships (Guerrero-Ramírez et al., 2017; Baert et al., 2018). We further expected (H4) the positive effect of tree diversity on soil microbial functions to be more pronounced in older experiments due to increased niche complementarity and stability (Reich et al., 2012; Guerrero-Ramírez et al., 2017).

## Materials & Methods

Soil samples were taken in 2013 from eleven tree diversity experiments that are part of the global network TreeDivNet (Verheyen et al., 2016; http://www.treedivnet.ugent.be/). Those experiments are independent with different experimental designs and plot configurations (Table 1). Experiments are distributed across four continents (Asia, Europe, North and South America) and four different biomes (boreal, temperate, tropical, subtropical; Olson et al., 2001), and differ in age with the youngest experiments running for three years and the oldest for fourteen years as of 2013 (i.e., the year of the sampling campaign; Fig. 1; Table 1). In total, 106 tree species were included in this study. Experiments have a mean number of diversity levels of 3.7 ± 1.0, with diversity levels reaching from monocultures to 16 tree species in sub/tropical regions.

**Table 1.** List of tree diversity experiments, which contributed to the study (alphabetical order). All experiments differ in their plot architecture as indicated by different number of diversity levels, and the gradient of diversity. Further, experiments differ in experimental age, number of sites and blocks, as well as in plot size, tree distance, tree density, and species pool (see Table S1). For the BEF-China and Sabah experiment, only a fraction of the whole diversity gradient was sampled (the respective missing richness levels are indicated by square brackets). The BIOTREE-FD experiment has only one species richness level (with four species per plot), but mixtures differ in their functional diversity (FD; indicated by a * in the Table). The number of plots only considers plots that enter the analysis, i.e., controls without trees were excluded as well missing measurements. The total number of existing plots is given in square brakets.

| Experi-<br>ment | Country | Biome | Age<br>(years) | Altitude<br>(m) | Former<br>land use | n<br>Sites | n<br>Blocks | n<br>Diversity<br>levels | Species<br>richness<br>levels | Plot size<br>(m<sup>2</sup>) | n Plots | Minimal<br>tree<br>distance<br>(m) | Tree<br>density<br>(trees m<sup>-2</sup>) | Reference |
| --- | --- | --- | --- | --- | --- | --- | --- | --- | --- | --- | --- | --- | --- | --- |
| Bangor | UK | temperate | 9 | 1 | forest | 1 | 2 | 3 | 1,2,3 | from 45 to<br>196 | 80 [92] | 1 | 1 | http://www.treedivnet.ugent.be/ExpBangor.html |
| BEF-China | China | subtropical | 4 | 190 | forest | 2 | NA | 5 | 1,2,4,8,16[,24] | 666.6 | 60 [566] | 1.29 | 0.6 | Bruehlheide et al., 2014;<br>doi: 10.1111/2041-210X.12126 |
| BIOTREE-FD | Germany | temperate | 10 | 400-415 | pasture | 1 | 4 | 4* | 4 (+FD) | 1700 | 24 [25] | 1 | 0.7 | Scherer-Lorenzen et al., 2007;<br>doi: 10.1016/j.ppees.2007.08.002 |
| FORBIO | Belgium | temperate | 3 | 398, 56, 13 | forest, arable<br>land | 3 | 6 | 4 | 1,2,3,4 | 1764 | 126<br>[127] | 1.5 | 0.4 | Verheyen et al., 2013;<br>doi: 10.5091/plecevo.2013.803 |
| IDENT<br>Auclair | Canada | temperate | 3 | 333 | pasture | 1 | 4 | 3 | 1,2,6 | 14.4 | 187<br>[192] | 0.4 | 8.2 | Tobner et al., 2013;<br>doi: 10.1007/s00442-013-2815-4 |
| IDENT<br>Cloquet | USA | temperate | 3 | 383 | forest | 1 | 4 | 3 | 1,2,6 | 14.4 | 190<br>[192] | 0.4 | 8.2 | Tobner et al., 2013<br>doi: 10.1007/s00442-013-2815-4 |
| Kreinitz | Germany | temperate | 8 | 115 | agricultural | 1 | 2 | 5 | 1,2,3,5,6 | 25 | 96 [98] | 0.8 | 1.2 | Hantsch et al., 2014;<br>doi: 10.1007/s00442-013-2815-4 |
| ORPHEE | France | temperate | 5 | 60 | forest | 1 | 2 [8] | 5 | 1,2,3,4,5 | 400 | 61 [256] | 2 | 0.3 | Castagneyrol et al., 2013;<br>doi: 10.1111/1365-2745.12055 |
| Sabah | Borneo | tropical | 11 | 102 | forest | 1 | NA [2] | 2 | 1,[4,]16 | 40000 | 27 [124] | 3 | 0 | Hector et al., 2011;<br>doi: 10.1098/rstb.2011.0094 |
| Sardinilla | Panama | tropical | 10 | 70 | forest | 2 | 6 | 3 | 1,2,5[,9,18] | 2025 | 46 [46] | 3 | 0.1 | Scherer-Lorenzen et al., 2007;<br>doi: 10.1111/j.2007.0030-1299.16065.x |
| Satakunta | Finland | boreal | 14 | 35 | forest | 3 | NA | 4 | 1,2,3,5 | 400 | 113<br>[163] | 1.5 | 0.4 | http://www.sataforestdiversity.org |
| **Average** |  |  | **7.2 ± 3.9** | **183.6 ±**<br>**163.8** |  |  |  | **3.7 ± 1.0** |  | **3937.5 ±**<br>**11381.2** | **total:**<br>**1010** | **1.6 ± 0.9** | **1.1 ± 1.2** |  |

**Figure 1:**
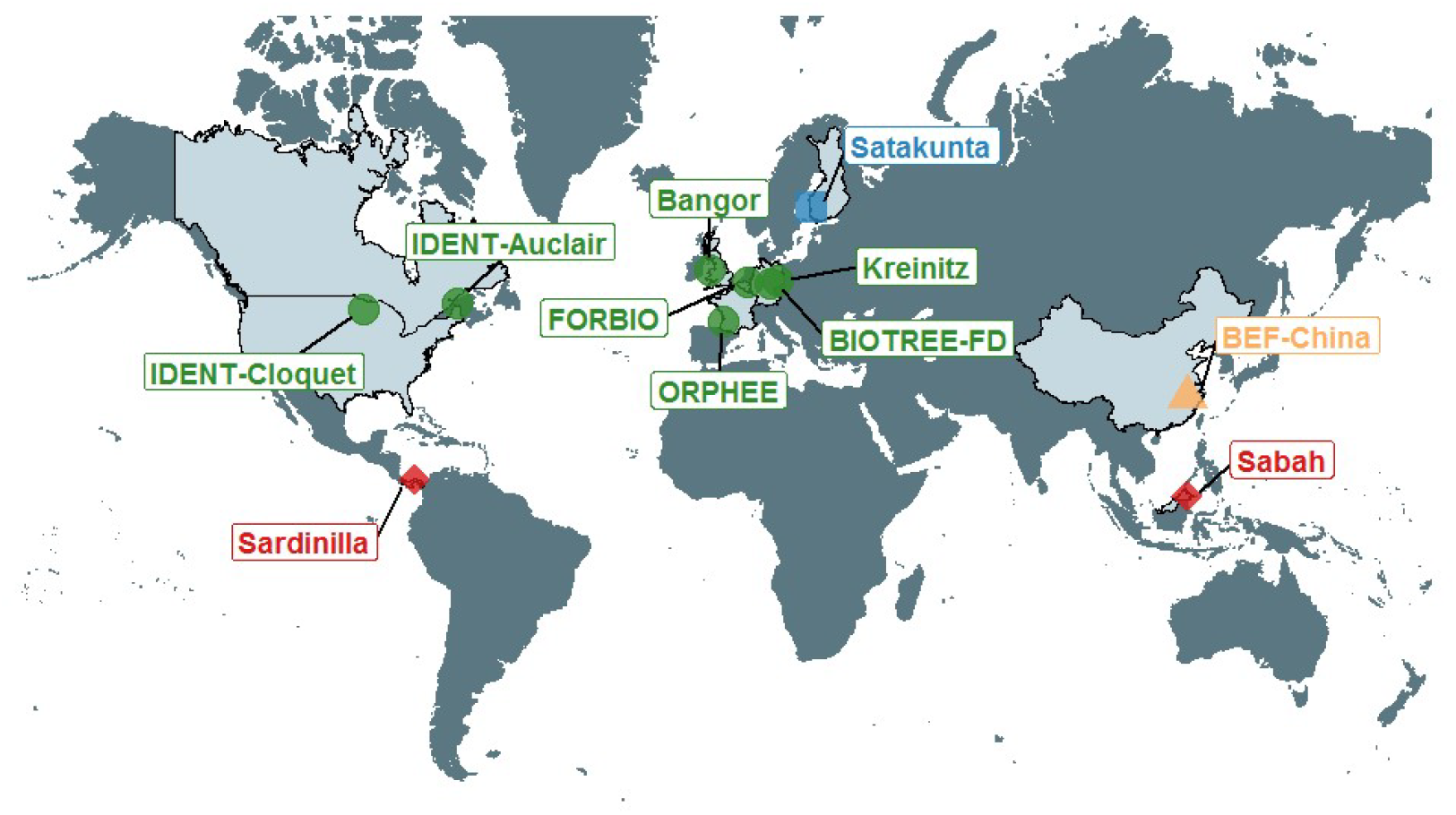
Locations of the eleven tree diversity experiments and assignment to biomes (Olsen et al 2001). Details of the locations and experimental designs are shown in Table 1.

### Soil sampling

Soil samples were taken from a depth of 0 – 10 cm, excluding the litter layer using a soil corer. Depending on the size of the experimental plot, different numbers of subsamples were taken per plot to create one composite sample. For plots <100 m^2^, three subsamples were taken per plot, while ten subsamples were taken for plots >100 m^2^. These subsamples were taken to capture spatial heterogeneity of the plot and to represent as many different combinations of tree species as possible. Immediately after sampling, soil samples were stored at 5°C until sieving at 2 mm, and then were stored afterwards at −20°C until shipping in the local laboratories to minimize changes in the microbial activity, biomass, and composition. Altogether, 1008 plots were sampled across the eleven tree diversity experiments in the framework of this study (Table 1).

### Measurement of soil microbial functions

Before the start of the microbial measurements, samples were kept at +20°C for five days to unfreeze and to adapt the soil microbial community to a constant and standardized temperature. Three different soil microbial community functions were assessed using an automated O_2_ micro-compensation system (Scheu, 1992). First, basal respiration (µl O_2_ h^−1^ g^−1^ dry soil) was measured as the mean oxygen consumption per hour without the addition of any substrate. The mean oxygen consumption was measured for hours 15 to 20. Basal respiration reflects the active part of the soil microbial community at the time of sampling. Second, microbial biomass C was measured by substrate-induced respiration, i.e., the respiratory response of microorganisms to glucose (and water) addition. To saturate catabolic microbial enzymes, 8 mg glucose g^-1^ soil dry weight was added as an aqueous solution to the soil samples. Substrate-induced respiration within the first 10 h was taken as the maximum initial respiratory response (MIRR) – a period where microbial growth has not started. Microbial biomass (µg C g^−1^ dry soil) was calculated as 38 × MIRR (µl O_2_ h^−1^ g^−1^ dry soil) according to Beck et al. (1997). By providing water and glucose, the maximum potential of the living microbial biomass is activated that is able to use glucose, whereas for basal respiration only a fraction of the entire community is active. Third, microbial specific respiratory quotient (µl O_2_ mg^−1^ Cmic h^−1^) was calculated as the ratio of basal respiration and soil microbial biomass.

The specific respiratory quotient is a measure of soil microbial carbon use efficiency. Carbon use efficiency is high when microbial biomass can be built up without high investment in respiration, which is indicated by a lower specific respiratory quotient. All measurements were conducted at 20°C in an air-conditioned laboratory using the same analytical devices (RMS Schuller, Darmstadt, Germany).

### Diversity metrics

The vast majority of TreeDivNet sites have an experimental gradient in tree species richness, which was used for our analyses, but there is also one experiment having a functional diversity gradient at a constant level of species richness only (BIOTREE-FD, see Table 1). In addition, we aimed at testing a tree diversity metric representing the functional diversity of the tree stands. Since comparable trait measurements are not available from all experiments, some tropical species are not present in trait databases, and because traits demonstrate plasticity to their abiotic and biotic environment (i.e. intraspecific trait variation), we chose not to use data from trait databases and to use phylogenetic diversity instead as a proxy for functional diversity. Phylogenetic diversity indices have been shown to be good predictors of biodiversity-ecosystem functioning relationships (e.g., Flynn et al., 2011; Craven et al., 2018) and are suggested to work when key functional traits are not available (Paquette *et al.*, 2015). We used the molecular phylogeny from previous studies (Zanne et *al.*, 2013, 2014) as a backbone to build a phylogeny of all species within the listed tree diversity experiments, conservatively binding species into the backbone using dating information from congeners in the tree. We used a set of different measures of phylogenetic diversity, i.e., MPD (mean phylogenetic diversity), MNTD (mean nearest taxonomic distance), and the standardized version of both to account for correlation with species richness. MPD was found to correlate less (using pearson correlation) with log species richness and was used in all following analyses as phylogenetic diversity metric (Table S1). Mean phylogenetic diversity (MPD) was calculated using the *comparative.comm* function in *pez* (Pearse *et al.*, 2015). Taxonomic names of tree species were standardized using the website http://tnrs.iplantcollaborative.org/index.html.

### Soil characteristics

We included a set of moderators in our models, which were shown to have an effect on soil microbial functions and reflect the designs and local conditions of the different experiments (Table 2). Gravimetric soil water content was measured as % H_2_O from fresh soil weight by drying the whole sample for three days at 75°C. Soil pH, soil C (%), soil nitrogen (soil N) (%), and soil C-to-N ratio (soil CN) were measured at the block level to obtain information about soil quality characteristics of each experiment. Therefore, equal proportions of dry soil were weighed from each sample to form a composite sample. The whole sample was ground, and a fraction of 10 g was used for pH measurements by adding 0.01 m CaCl_2_. Soil C and N concentrations were analyzed by using the ground soil with an elemental analyzer (Vario EL Cube, Elementar). We further extracted clay (%), sand (%), and silt (%) content from the SoilGRIDS database (Hengl *et al.*, 2014).

**Table 2:** Akaike information criterion (AIC) after model comparison for models using log tree species richness (SR) or mean phylogenetic distance (MPD) to evaluate their explanatory power for soil basal respiration, microbial biomass, and the respiratory quotient. Models with lower AICc have a better model fit. df: degrees of freedom. Marginal R^2^ represents the variance explained by fixed factors, whereas conditional R^2^ represents the variance explained by both fixed and random factors. Bold values indicate the best model fit for each response variable.

| Response variable | Biodiversity metric | df | AIC | R2 fixed (marginal) | R2 random (conditional) |
| --- | --- | --- | --- | --- | --- |
| Basal respiration | log SR | 20 | 1159.25 | 0.52 | 0.84 |
|  | MPD | 21 | 1164.89 | 0.53 | 0.84 |
| Microbial biomass | log SR | 21 | 892.69 | 0.50 | 0.87 |
|  | MPD | 21 | 892.53 | 0.49 | 0.88 |
| Respiratory quotient | log SR | 21 | 1688.96 | 0.42 | 0.74 |
|  | MPD | 21 | 1695.69 | 0.41 | 0.74 |

### Environmental conditions

For each experimental site, we extracted mean annual temperature (MAT), annual precipitation (MAP), potential evapotranspiration (PET), and aridity index from the WorldClim database (http://www.worldclim.org/current) with 2.5 arc-minutes resolution. In addition, we obtained the age of the experiment (years), altitude (m), tree distance (m), and tree density (trees m^−2^) from publications associated with each experiment (Table 1) and the TreeDivNet website (http://www.treedivnet.ugent.be/). Biomes were assigned based on Olson *et al.* (2001).

### Data analysis

Prior to analysis, all data were centred and scaled using the scale function from the *base* package in R, and the distributions of response variables were checked visually (Zuur *et al.*, 2010). To minimize the effects of collinearity, an automatic stepwise selection of the variables using VIF (variation inflation factor) was applied (https://gist.github.com/fawda123/4717702). The function calculates the VIF of all explanatory variables and removes the variable with the highest VIF. The procedure starts again with the reduced set of explanatory variables until all VIF values are below the threshold of 3 (Zuur *et al.*, 2010). The final list of variables were aridity, experimental age, PET, soil pH, soil CN, soil N, and tree density that were included as fixed factors in the models. In addition to VIF, correlations of all explanatory variables were calculated as Spearman’s rho using the *cor* function from the stats package (Table S2).

For each response variable (basal respiration, microbial biomass, and respiratory quotient), we fitted two separate linear mixed-effects models that included either log species richness or phylogenetic diversity as a measure of tree diversity. Random terms were block nested in site, and site nested within experiment (see Table 1 for information about levels of the random effects). We used *optimx* algorithm to ensure model convergence. As three-way interactions of the fixed factors result in extremely high variance inflation (VIF), we fitted models with two-way interactions only. Interactions with VIF >3 were removed from the full model (Zuur *et al.*, 2010). After model reduction, model selection based on AICc was used to evaluate if models with species richness (log-scale) or mean phylogenetic distance were more parsimonious. The Kenward-Rogers approximation was used to test for the significance of fixed effects and degrees of freedom. Marginal and conditional R^2^ were calculated using the function *r.squaredGLMM* from the *MuMIn* package. Marginal R^2^ represents the variance explained by the fixed effects, whereas conditional R^2^ represents the variance explained by both fixed and random effects. We checked model assumptions of the most parsimonious models by fitting model residuals versus fitted models. Basal respiration and the respiratory quotient were log-transformed to achieve the requirements of parametric statistical tests. Model fits of the mixed effects models were used to plot estimates using the function *plot_model* from the package *sjPlot*. Significant interactions were plotted using *ggpredict* from the package *ggeffects*. To do so, continuous variables were divided into constant levels automatically. All statistical analyses were performed in R (version 3.3.5) (R Core Team, 2016).

## Results

We used eleven tree experiments with 106 different tree species distributed across four biomes to test tree diversity-soil microbial functioning relationships. Mean soil basal respiration (± SD) was 2.06 ± 1.94 µl O_2_ h^−1^ g soil dw^−1^, with the lowest values in the FORBIO experiment in Belgium (min: 0.08 µl O_2_ h^−1^ g soil dw^−1^) and highest values in the boreal Sastakunta experiment in Finland (max: 15.26 µl O_2_ h^−1^ g soil dw^−1^) (Fig. 2). Similarly, we found lowest soil microbial biomass values in the FORBIO experiment (min: 11.85 µg Cmic g soil dw^−1^) and highest values in the Satakunta experiment (max: 2501.54 µg Cmic g soil dw^−1^). Mean microbial biomass was 435.51 ± 325.03 µg Cmic g soil dw^−1^. The respiratory quotient was lowest (i.e., high carbon use efficiency) in BIOTREE-FD in Germany (max: 0.008 µl O_2_ µg^−1^ Cmic h^−1^), but most of the low values were measured in both IDENT experiments. The highest respiratory quotient was measured in ORPHEE in France (min: 0.0395 µl O_2_ µg^−1^ Cmic h^−1^), whereas FORBIO and Satakunta also showed very low values. The grand mean across experiments was 0.0052 ± 0.0031 µl O_2_ µg^−1^ Cmic h^−1^. Mean soil water content was 17.2 ± 11.5%, and the driest soil was found in IDENT Cloquet in Minnesota, USA (min: <0.1%), whereas highest values were measured in Satakunta (max: 58.5%).

**Figure 2:**
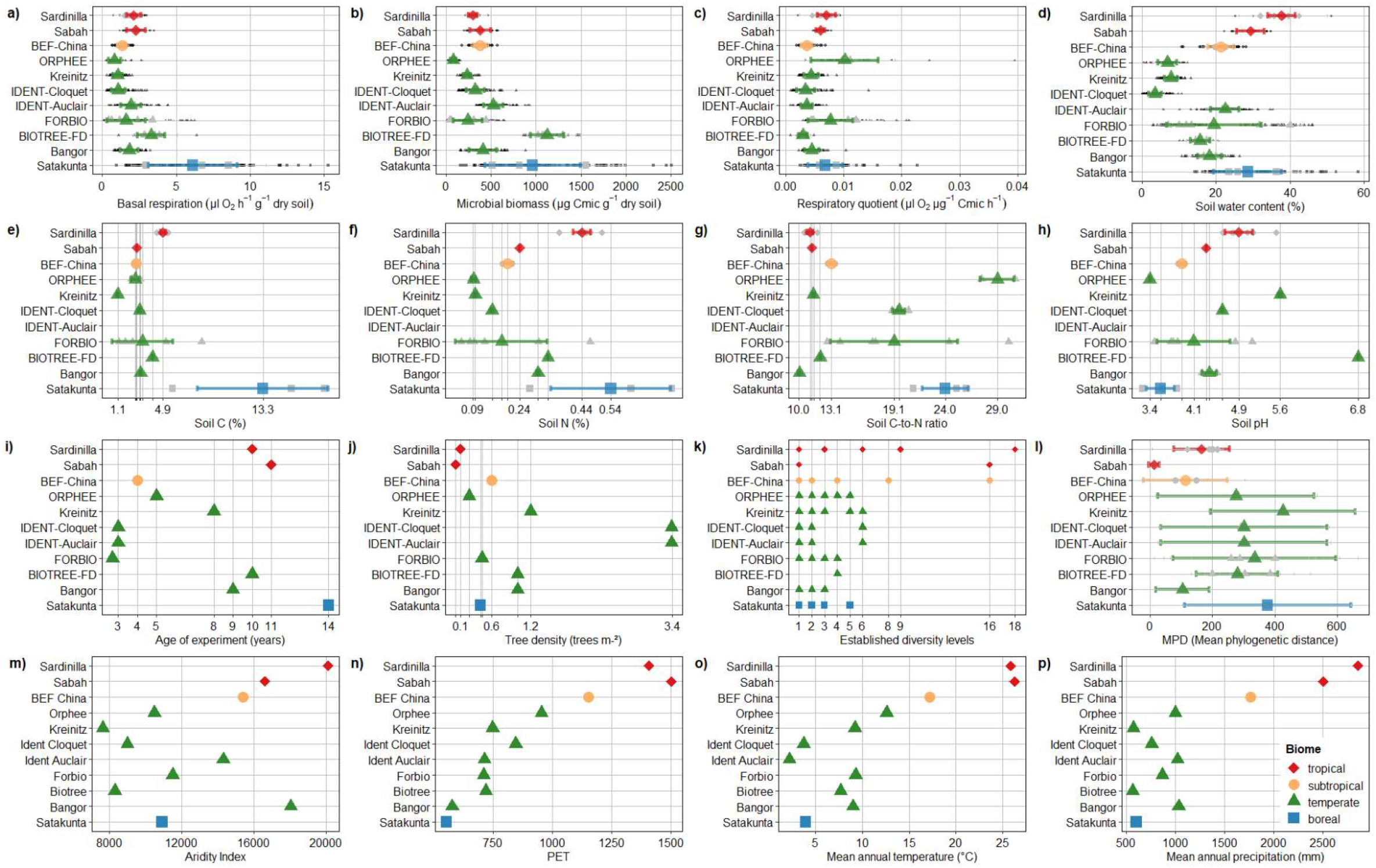
Overview of measured response variables [a-d] and predictors [e-p] across global tree diversity experiments. Grey symbols and error bars (SD) refer to blocks or sites, whereas colored symbols refer to the grand mean ± SE of each experiment. Predictors for describing soil quality [e-h] were measured at the block level (if present, otherwise site level), and, therefore, values of single blocks or sites (grey symbols) have no variance. Small black symbols in panel a-d) represent single measurements. Colors and symbols represent the four different biomes (see legend). Vertical lines with irregular distance indicate specific measurements. Therefore, numbers on the x-axis are only presented when values do not overlap.

### H1: Tree diversity increases soil microbial functions

Overall, tree diversity did not significantly increase basal respiration, microbial biomass, or decreased the respiratory quotient (Fig. 3). However, for basal respiration, tree diversity had weak but statistically significantly interaction effects with PET, soil CN, and tree density, with a positive effect of tree species richness at medium and low PET levels (Fig. 4a) and low CN levels (Fig. 4b). Moreover, increasing MPD had a positive effect on basal respiration when tree density was high (Fig. 4c). Soil microbial biomass and the respiratory quotient were not significantly influenced by any interaction with tree diversity (Fig. 3).

**Figure 3:**
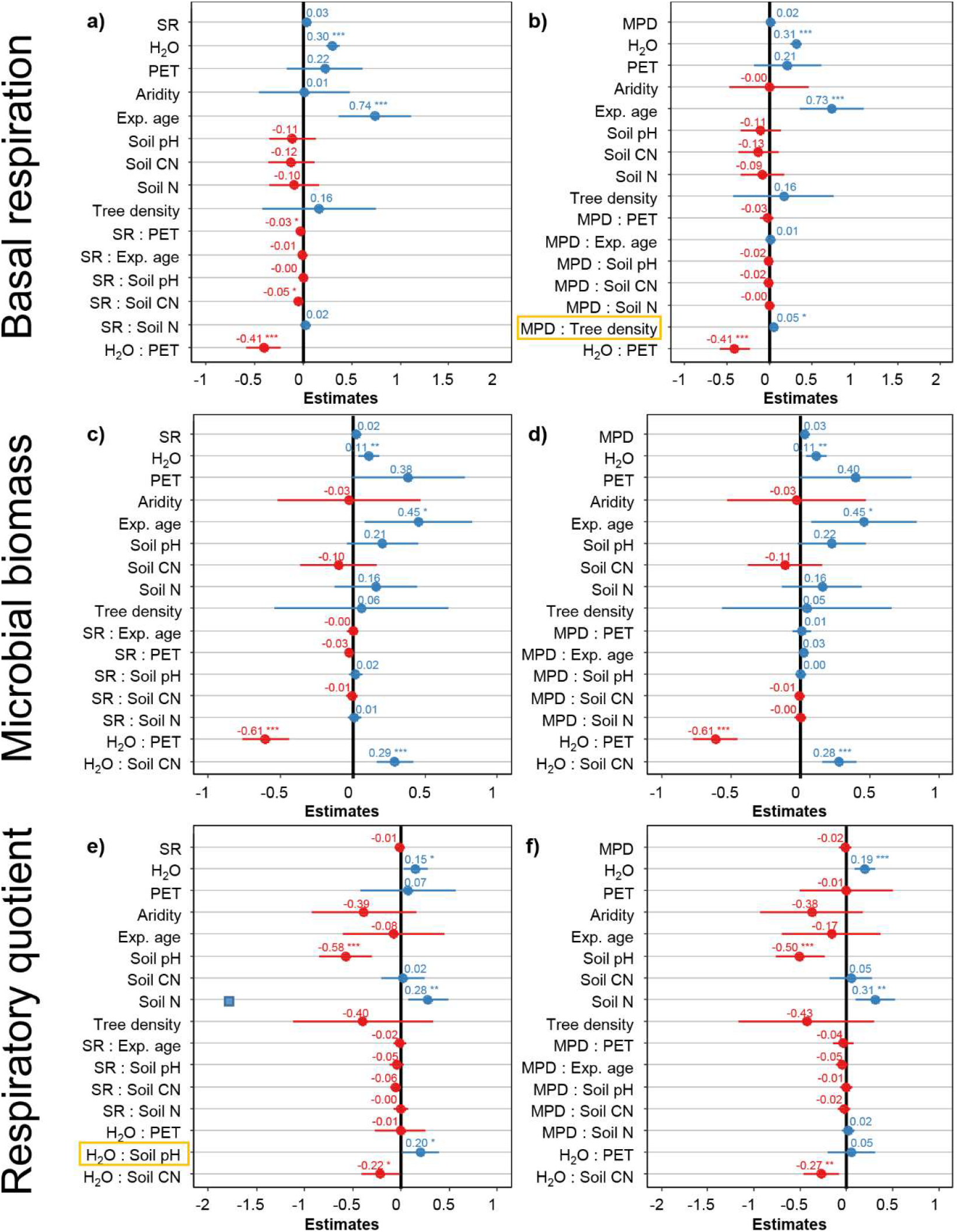
Coefficient estimates for three soil microbial functions (basal respiration, microbial biomass and the respiratory quotient) as affected either by log species richness (SR; panels a,c,e) or mean phylogenetic distance (MPD; panels b,d,f) and a set of biotic and abiotic variables that remained after accounting for collinearity. Blue colors indicate a positive trend of the fixed factor on the response variable, whereas red indicates a negative relationship. Yellow boxes highlight the only additional interactions between the two models differing in the tree diversity metric. H_2_O: soil water content, PET: Potential evapotranspiration, Exp. age: Experimental age. Asterisks indicate strength of the significance level with ***: P<0.001; **: P< 0.01; *: P<0.05. Detailed information about significant interactions is given in Fig. 4.

**Figure 4:**
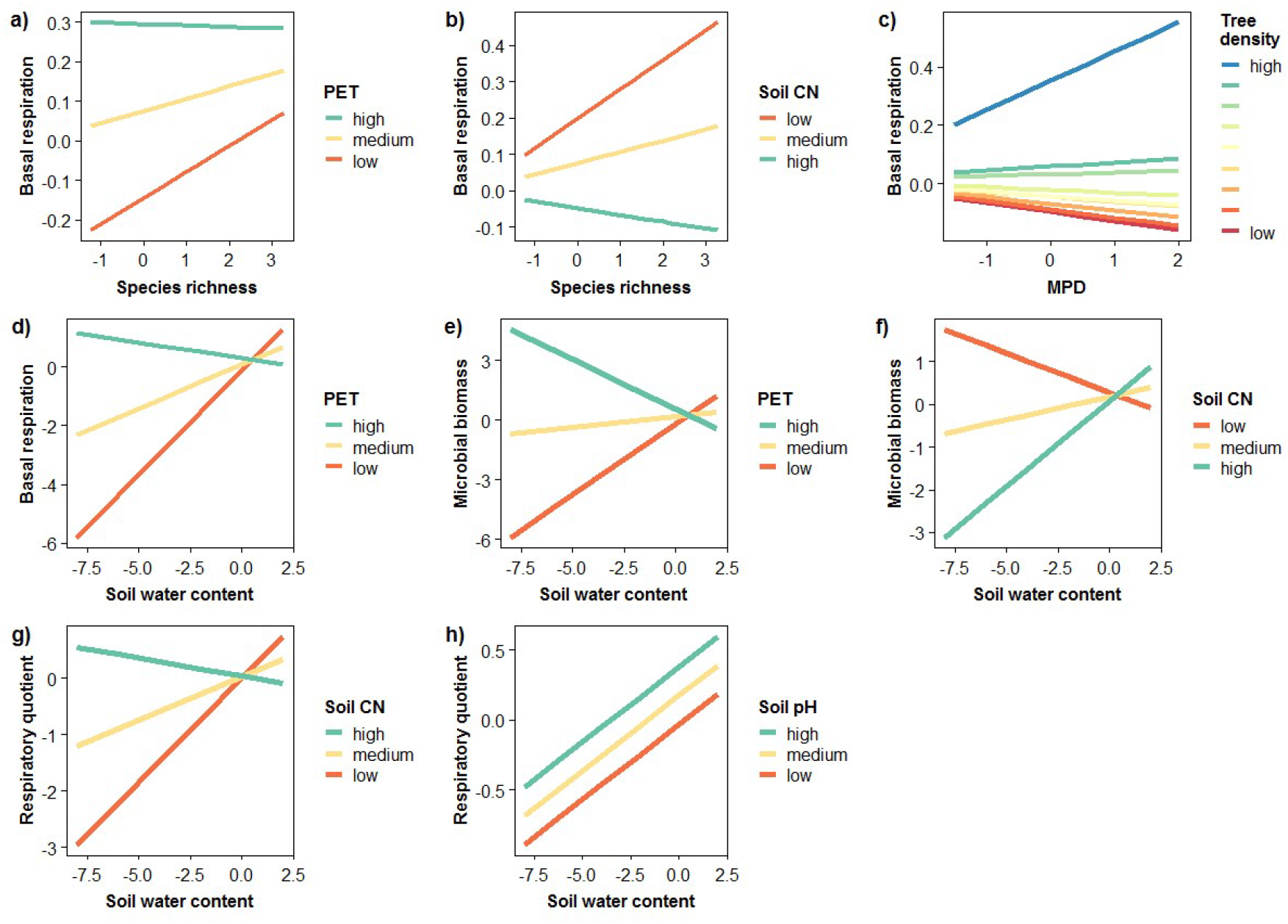
Predicted interaction plots of significant interactions showing the relationship for the three microbial functions: basal respiration (a-d; µl O_2_ h^−1^ g soil dw^−1^), microbial biomass (e-f; µg C_mic_ g soil dw^−1^), and the respiratory quotient (g-h; µl O_2_ µg^−1^ C_mic_ h^−1^) based on linear mixed models. Continuous variables were automatically subdivided into levels.

### H2: The effect of tree diversity on soil microbial functions is higher for phylogenetic diversity than species richness

Our models were able to explain 41% to 53% (R2 marginal, fixed factors) of the variation and even more when accounting for the random effects (74% to 88%; Table 2). For all single factor effects (SR, H_2_O, PET, Aridity index, Experimental age, soil pH, Soil CN, Soil N, and tree density), models using species richness as a diversity metrics did not differ from models using MPD as indicated by very similar estimates, AICs, and R^2^ (Fig. 3; Table 2). Similarly, all but two interactions (yellow boxes in Fig. 3) were the same in the models of the two diversity metrics. Differences occurred for basal respiration, here, MPD showed a significant interaction with tree density, which did not occur in the species richness model. Second, for respiratory quotient, soil water content significantly interacted with soil pH, an interaction that did not appear in MPD models.

### H3: High soil C content, soil water content, and soil pH increase soil microbial functions

Soil C was not retained in the final models, but showed a highly significant correlation with soil N (Table 2) which stayed in the models. Generally, soil N significantly increased the respiratory quotient indicating lower carbon-use efficiency with increasing N levels (Fig. 3e,f). Soil water content increased all soil microbial functions significantly. Soil water content was involved in many significant interactions, indicating that this variable modulated effects of many other factors. All three soil microbial functions were, however, affected by a different set of interactions. Basal respiration increased strongest with soil water content when PET was low (Fig. 4d). In contrast, at high PET soil basal respiration slightly decreased with increasing soil water content. However, at high soil water content, basal respiration was overall highest and the different levels of PET were of low importance. As for basal respiration, the importance of PET and soil CN for soil microbial biomass was reduced at high soil water content levels (Fig. 4e,f). High PET and low CN reduced soil microbial biomass with increasing soil water content, whereas low PET and high soil CN increased it. The main pattern, that high soil water content overwhelmed the importance of a second driver, was also found for the respiratory quotient and soil CN. At low soil water content, soil CN had the strongest effect on respiratory quotient, with the highest values at high soil CN, while increasing soil water content rendered the effect of soil CN non-significant (Fig. 4g). High soil pH gradually increased the respiratory quotient (lower carbon use efficiency) compared to medium and low soil pH with increasing soil water content (Fig. 4h).

### H4: Positive BEF relationships are more pronounced in older tree diversity experiments

Older experiments had significantly higher basal respiration and soil microbial biomass, but this positive effect was not strengthened by tree diversity or any other environmental factor (Fig. 2).

## Discussion

We tested abiotic and biotic drivers of soil microbial functions in eleven tree diversity experiments in a variety of different contexts across the globe and found soil microbial functions to be dominated by abiotic rather than by biotic factors. However, effects of tree diversity on soil microbial respiration were highly context dependent and were strongest where PET and soil CN were low and tree density was high.

Our first hypothesis suggested positive effects of tree diversity on soil microbial functions, but we did not find any significant main effect. However, tree species richness and MPD depended on other environmental variables in affecting basal respiration. By contrast, soil microbial biomass and the respiratory quotient were not affected by tree diversity at all. None or weak tree diversity effects on soil microorganisms have been shown in many previous studies, highlighting that tree species identity may be a more important driver of soil microorganisms and soil functions (e.g., Khlifa et al., 2017; Gottschall et al., 2019). Our study suggests that soil microorganisms are mainly influenced by abiotic drivers, which are also important in modulating tree diversity effects. However, tree species with strong impact on soil characteristics like soil pH, e.g. evergreen coniferous species, may explain more variation (Reich et al., 2005). For instance, single tree species can affect the structure of the litter layer with significant influence on microclimatic conditions that drive soil microbial functions (Gottschall et al., 2019). Therefore, improved data on litter and root traits as well as their influence on soil quality and microclimate are required to improve the mechanistic understanding of tree effects on soil functions (Laliberté, 2017). Using belowground traits, rather than aboveground traits, is essential to predict soil functions, especially as different mechanisms are expected to operate belowground. For instance, aboveground traits for nutrient acquisition converge at high P-use efficiency, whereas belowground traits of nutrient acquisition strategies increase (Zemunik et al., 2015). The growing network of global tree diversity experiments (Verheyen et al., 2016; Paquette et al., 2018), and other similar networks (Borer et al., 2014) will allow for coordinated approaches and should aim to directly measure belowground traits to identify abiotic and biotic drivers of soil microbial functions.

Our second hypothesis suggested stronger effects of phylogenetic diversity than taxonomic diversity. Differences between our two models that included mean phylogenetic distance or tree species richness as predictors were small, although some other studies on plant biomass production in grassland provided stronger support that phylogenetic diversity is a better predictor of ecosystem functioning than species richness (Cadotte et al., 2008; but see e.g. Flynn et al., 2011). Microbial functions are mainly affected by abiotic drivers (Serna-Chavez et al., 2013), a finding that is also supported by the present study, thereby possibly masking effects of tree species richness and phylogenetic diversity. However, as tree diversity is a weak predictor of soil microbial functions in forests, trait-based approaches may provide more mechanistic insights.

We further hypothesized soil C, soil water content, and soil pH to be important drivers of soil microbial functions. Confirming this expectation, soil water content was the dominant driver affecting all soil microbial functions and interacting with all mentioned abiotic drivers. The high importance of soil water content for soil microbial processes has been shown in many studies (e.g., Schimel, 2018). The present study further emphasises that optimal water levels can attenuate the impact of other factors. Moderate levels of soil water content (i.e., between 50 – 70% of field capacity) generally increase microbial biomass and activity, whereas lower and higher levels reduce oxygen availability (Franzluebbers, 1999), with negative impacts on soil microbial biomass and activity. We found that at high water levels, changes in soil pH, and soil CN had minor effects on the overall high values of soil microbial functions, suggesting that soil water availability is the principal driver. For instance, positive effects of high temperature on soil biological activity can only be achieved when soil water is not limited (Thakur et al., 2018), and nutrient availability can be increased by higher soil moisture via increasing diffusion of soluble organic substrates (Hungate et al., 2007). This suggests that sufficient soil water availability can mitigate other unfavorable abiotic effects, thereby increasing soil ecosystem functioning. On the other hand, tree communities subjected to dry conditions will be affected to a greater extent by soil characteristics without the buffering capacity of water. Therefore, to maintain soil ecosystem functioning, especially when faced with more frequent dry periods due to global change, tree species that use water more efficiently could be selected or communities that have a higher diversity in hydraulic traits may be better able to mediate ecosystem resilience to low soil water content (Anderegg et al., 2018, 2019). In addition, alternative strategies (e.g., leaving leaf litter on ground, applying mulch, planting a cover crop) are needed to enhance soil water content, which can be developed in parallel to the establishment of high diversity forests.

We did not find support for a strengthened biodiversity effect over time, although we found that experimental age increased soil basal respiration and microbial biomass significantly. Continuous inputs of resources and changes in the quality of these inputs over time increases the microbial pool and switches microbial communities to fungal dominance (Cline & Zak, 2015). We found that experimental age was significantly correlated with soil C (rho = 0.58, P = <.0001) and soil N (rho = 0.64, P = <.0001) suggesting that experimental age increases nutrient content and maybe soil C storage, potentially due to decreased CO_2_ loss by fungal communities (Malik et al., 2016). Following this argument, the increase in basal respiration, i.e., microbial activity, is suggested to increase carbon sequestration by adding more soil microbial necromass, i.e., dead microbial biomass, into the soil carbon pool (Schmidt et al., 2011; Lange et al., 2015). Moreover, microbial necromass was shown to be a stronger driver of soil organic carbon than plant-derived substrates in grassland soils across the globe (Ma et al., 2018). The accumulation of soil microbial necromass and, thereby, increasing soil organic carbon levels, has been linked to complex interactions between climate (soil moisture/aridity) and soil texture (mainly driven by clay content), and further interactions with nutrient content and substrate quality (Ma et al., 2018). Soils with a higher soil organic matter content may have a higher soil water retention capacity than soils with higher proportion of mineral components. Further, IPCC suggests to increase global forest areas to capture CO_2_ to keep warming below 1.5°C (IPCC, 2018). This equals to ~24 million hectares of forest every year until 2030 (Lewis & Wheeler, 2019). Although many countries are committed to plant and restore forests or to promote more natural forests with a higher degree of functions, recent analyses show that those activities involve commercial forests with a lower ability to store carbon than more natural forests with a higher degree of multifunctionality (Lewis & Wheeler, 2019). Exploring interactions between abiotic and biotic factors in driving soil microbial functions and carbon storage in future studies is pivotal in order to get a more mechanistic understanding of driving forces of soil carbon storage.

## Conclusion

Global analyses of biodiversity-ecosystem functioning relationships aim to identify general patterns, context-dependencies, and underlying mechanisms to predict and mitigate the consequences of species loss for human well-being. Results of the present study indicate that higher soil water levels increase soil microbial functions, which can have important consequences for soil carbon dynamics and sequestration. Moreover, improved data on litter and belowground plant traits as well as soil quality are required to improve the mechanistic understanding of tree effects on soil processes. Notably, results of tree diversity experiments may have important practical implications for ecological applications as many degraded ecosystems are reforested, and recommendations regarding how to enhance the multifunctionality of these reforestation efforts are urgently needed. Our data suggest to aim for alternative strategies to maintain soil water content to maintain high soil microbial functions in a changing climate. Context-dependent tree diversity effects further indicate that management recommendations have to consider local environmental conditions.

## Acknowledgements

German Centre for Integrative Biodiversity Research (iDiv) Halle-Jena-Leipzig funded by the German Research Foundation (DFG FZT 118). We thank Alfred Lochner, Anja Zeuner and Silke Schroeckh for measuring soil abiotic variables and support with respiration measurements.

BEF-China was funded by German Research Foundation (DFG FOR 891/1-3).

The BIOTREE experiment in Bechstedt has been established by the Max-Planck-Institute for Biogeochemistry Jena, Germany, and we are grateful to Prof. Dr. Ernst-Detlef Schulze for initiating and supporting this project. BIOTREE receives basic funding through the Chair of Geobotany, Faculty of Biology, University of Freiburg. The BIOTREE site Bechstedt is maintained by the Federal Forestry Office Thüringer Wald (Bundesforstamt Thüringer Wald).

FORBIO was partly supported by the Walloon forest service (SPW-DNF), through the 5-year ‘Accord-cadre de recherche et de vulgarisation forestières’ programme

The Kreinitz Experiment has been funded by the Helmholtz Centre for Environmental Research—UFZ. We are grateful to the many colleagues who have assisted with the establishment and maintenance of the experiment and who are too numerous to be listed. In particular, we acknowledge the Departments of Community Ecology, Soil Ecology, Soil System Science and Environmental Microbiology, and the team of the Bad Lauchstädt field station of the UFZ.

Sardinilla has been mainly managed by José Monteza the site manager, Lady Mancilla, and their fieldworkers. Support came also from the Smithsonian Tropical Research Institute, Panama.

The Satakunta Experiment has been established by the University of Turku with the funding from the Academy of Finland. We are grateful to Dr Kai Ruohomäki from the Department of Biology, University of Turku for help with the establishment and maintenance of the experiment.

## Supplementary material

**Table S1:**
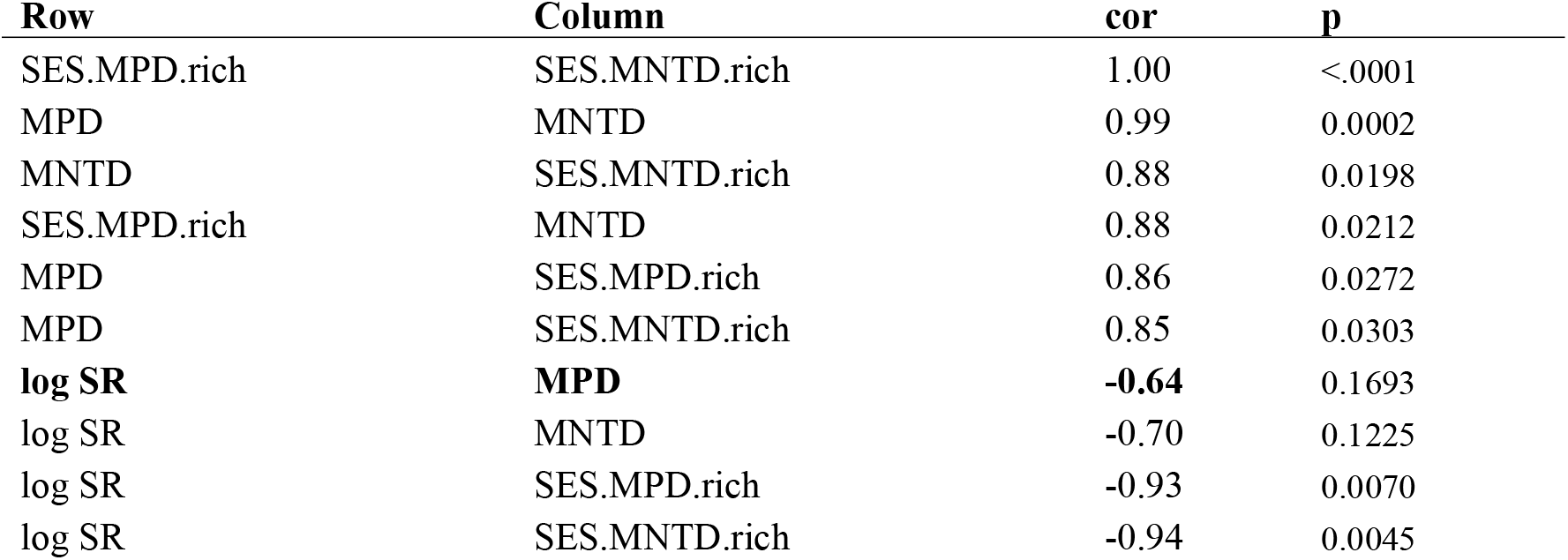
Correlation of diversity metrics using pearson correlation (cor). MPD: Mean phylogenetic diversity, MNTD: mean nearest taxon distance, log SR: log of species richness, SES indicates the standardized version of MPD and MNTD. The phylogentic diversity metric that correlates less with log species richness is marked in bold and is used for further analyses.

| Row | Column | cor | p |
| --- | --- | --- | --- |
| SES.MPD.rich | SES.MNTD.rich | 1.00 | <.0001 |
| MPD | MNTD | 0.99 | 0.0002 |
| MNTD | SES.MNTD.rich | 0.88 | 0.0198 |
| SES.MPD.rich | MNTD | 0.88 | 0.0212 |
| MPD | SES.MPD.rich | 0.86 | 0.0272 |
| MPD | SES.MNTD.rich | 0.85 | 0.0303 |
| **log SR** | **MPD** | **-0.64** | 0.1693 |
| log SR | MNTD | -0.70 | 0.1225 |
| log SR | SES.MPD.rich | -0.93 | 0.0070 |
| log SR | SES.MNTD.rich | -0.94 | 0.0045 |

**Table S2:** List of scaled explanatory variables to be included as fixed factors in the models and their correlation (Spearman’s rho and significane level). Source indicates if values were measured on the plot (Measurement) or block (Block) level, retrieved from a specific database or just reflect experimental conditions (Experiment). # indicates if the listed variables stayed in the final mixed effect model after inspecting colinearity using VIF. H_2_O = soil water content, MAP = Mean annual precipitation, MAT = Mean annual temperature, Aridity = Aridity Index, PET = Potential evapotranspiration, Exp. age = Experimental age, T density = Tree density, T distance = Tree distance, log SR = log Species richness, MPD = mean phylogenetic distance.

|  |  | Soil characteristics |  |  |  |  |  |  |  | Environmental conditions |  |  |  | Experimental condition |  |  |  | Diversity metric |  |
| --- | --- | --- | --- | --- | --- | --- | --- | --- | --- | --- | --- | --- | --- | --- | --- | --- | --- | --- | --- |
| Source |  | H2O<sup>#</sup> | Soil pH<sup>#</sup> | Soil C | Soil CN<sup>#</sup> | Soil N | Sand | Silt | Clay | MAP | MAT | Aridity<sup>#</sup> | PET<sup>#</sup> | Exp. age<sup>#</sup> | T density<sup>#</sup> | T distance | Altitude | log SR<sup>#</sup> | MPD<sup>#</sup> |
| Measurement | H2O<sup>#</sup> | 1 |  |  |  |  |  |  |  |  |  |  |  |  |  |  |  |  |  |
| Block | Soil pH<sup>#</sup> | -0.27*** | 1 |  |  |  |  |  |  |  |  |  |  |  |  |  |  |  |  |
| Block | Soil C | 0.43*** | -0.4*** | 1 |  |  |  |  |  |  |  |  |  |  |  |  |  |  |  |
| Block | Soil CN<sup>#</sup> | -0.2*** | -0.45*** | 0.41*** | 1 |  |  |  |  |  |  |  |  |  |  |  |  |  |  |
| Block | Soil N | 0.56*** | -0.28*** | 0.91*** | 0.09* | 1 |  |  |  |  |  |  |  |  |  |  |  |  |  |
| SoilGRIDS | Sand | -0.12** | -0.22*** | 0.02ns | 0.36*** | -0.18*** | 1 |  |  |  |  |  |  |  |  |  |  |  |  |
| SoilGRIDS | Silt | -0.12*** | 0.28*** | -0.02ns | -0.3*** | 0.11** | -0.91*** | 1 |  |  |  |  |  |  |  |  |  |  |  |
| SoilGRIDS | Clay | 0.35*** | 0.1** | -0.02ns | -0.35*** | 0.23*** | -0.89*** | 0.61*** | 1 |  |  |  |  |  |  |  |  |  |  |
| WorldClim | MAP | 0.5*** | -0.09* | -0.14*** | -0.39*** | 0.12** | -0.49*** | 0.12*** | 0.79*** | 1 |  |  |  |  |  |  |  |  |  |
| WorldClim | MAT | 0.43*** | 0.03ns | -0.24*** | -0.39*** | 0.01ns | -0.38*** | -0.01ns | 0.73*** | 0.91*** | 1 |  |  |  |  |  |  |  |  |
| CGIAR-CSI | Aridity<sup>#</sup> | 0.57*** | -0.23*** | 0.02ns | -0.41*** | 0.33*** | -0.41*** | 0.15*** | 0.61*** | 0.78*** | 0.68*** | 1 |  |  |  |  |  |  |  |
| CGIAR-CSI | PET<sup>#</sup> | 0.15*** | 0.06ns | -0.32*** | -0.23*** | -0.17*** | -0.48*** | 0.16*** | 0.73*** | 0.86*** | 0.82*** | 0.39*** | 1 |  |  |  |  |  |  |
| Experiment | Exp. age<sup>#</sup> | 0.55*** | -0.02ns | 0.58*** | -0.05ns | 0.64*** | -0.3*** | 0.2*** | 0.35*** | 0.07* | 0.1** | 0.25*** | -0.18*** | 1 |  |  |  |  |  |
| Experiment | T density<sup>#</sup> | -0.64*** | 0.28*** | -0.25*** | 0.06ns | -0.34*** | -0.07* | 0.33*** | -0.23*** | -0.35*** | -0.57*** | -0.47*** | -0.08* | -0.49*** | 1 |  |  |  |  |
| Experiment | T distance | 0.63*** | -0.15*** | 0.18*** | -0.04ns | 0.34*** | -0.18*** | -0.21*** | 0.57*** | 0.69*** | 0.78*** | 0.54*** | 0.52*** | 0.42*** | -0.78*** | 1 |  |  |  |
| Experiment | Altitude | -0.32*** | 0.26*** | -0.31*** | -0.07* | -0.38*** | -0.08* | 0.24*** | -0.12** | -0.14*** | -0.35*** | -0.45*** | 0.12** | -0.56*** | 0.75*** | -0.46*** | 1 |  |  |
| Diversity metric | log SR<sup>#</sup> | 0.09** | 0.17*** | -0.03ns | -0.08* | 0.04ns | -0.1** | -0.02ns | 0.2*** | 0.2*** | 0.25*** | 0.07* | 0.21*** | 0.1** | -0.14*** | 0.29*** | -0.05ns | 1 |  |
| Diversity metric | MPD<sup>#</sup> | -0.15*** | 0.03ns | 0.11** | 0.18*** | -0.01ns | 0.29*** | -0.2*** | -0.32*** | -0.32*** | -0.27*** | -0.37*** | -0.24*** | -0.01ns | 0.06ns | -0.14*** | 0.05ns | 0.42*** | 1 |

\*\*\* = P < 0.0001; \*\* = P < 0.001; \* = P < 0.05; ns = P > 0.05

